# Lymphangiogenesis in Abdominal Aortic Aneurysm regulates the balance between resident and circulating eosinophils via a 15-lipoxygenase-dependent mechanism

**DOI:** 10.1101/2025.07.16.664018

**Authors:** A. Hostalrich, L. Verdu, E. Balzan, A. Bizos, M. Dubourdeau, B. Lebas, J. Malloizel-Delaunay, F. Morfoisse, E. Lacazette, AC. Prats, X. Chaufour, B. Garmy-Susini

## Abstract

The abdominal aortic aneurysm (AAA) is a chronic degeneration of the aortic wall involving an inflammatory response, aberrant remodeling of the extracellular matrix, and the development of microvessels. Among these, lymphatic capillaries develop in the adventitia. However, the role of lymphangiogenesis in AAA remains unclear.

Here, we confirmed the development of lymphatic vessels in both human and mouse AAA. This was associated with a decrease in specialized pro-resolving mediators (SPM) generated by 15-Lipoxygenase (15LO), an enzyme that controls resolution of inflammation in lymphatic diseases. Lymphatic selective depletion of 15LO (Prox1cre; Alox15^fl/fl^ mice) increased both systemic and resident eosinophil (EOS) accumulation in lesions, but had no effect on other immune cell populations. Mechanistically, *in vitro* depletion of 15LO in lymphatic endothelial cells (LEC) significantly decreased EOS adhesion. In contrast, 15LO LEC depletion improved transendothelial migration. *In vivo*, the rescue of 15LO using lentivector transduction modified the balance between resident and systemic EOS in favor of the resident ones and reduced the AAA lesion.

Altogether, we show that lymphatic vessels play a protective role in AAA by regulating the trafficking of EOS into the aorta wall.

## Introduction

Abdominal aortic aneurysm (AAA) is a complex vascular disease characterized by the progressive dilation of the abdominal aorta, resulting in a weakened vessel wall prone to rupture^1^. This condition is associated with significant morbidity and mortality, primarily due to the risk of aortic rupture, which often leads to fatal hemorrhage^2^. When patients develop an AAA larger than 5 and 5.5cm in maximum diameter in women and men respectively, surgery remains the only option as for these larger aneurysms the risk of rupture is usually higher than the risk of surgery^3,4^. To date, only surgery can treat AAA. Because of its asymptomatic nature, its precise prevalence is difficult to assess. Estimates vary between 1 and 5% of the population over 65 with tobacco as the main risk factor^5,6^. In 2025, AAA rupture still leads to a mortality rate of 80%^7^. While the exact mechanisms driving AAA formation and progression are not fully understood, chronic inflammation, immune dysregulation, and extracellular matrix degradation have emerged as central pathological features of the disease^8,9,10^. Recently, the lymphatic system, along with specific immune cell populations such as eosinophils, has gained attention for its role in modulating vascular inflammation and influencing AAA progression^11,12^. Additionally, the resolution of inflammation is recognized as a critical component in preventing the perpetuation of vascular damage in AAA^13,14^.

The lymphatic system plays an essential role in maintaining fluid balance, immune surveillance, and the resolution of inflammation^15,16^. It facilitates the clearance of interstitial fluid, immune cells, and inflammatory mediators from tissues, helping to regulate the immune response and limit chronic inflammation^17^. In the context of AAA, impaired lymphatic function and disrupted lymphangiogenesis have been observed, which may exacerbate local inflammation and contribute to vessel wall degeneration^11^. Defective lymphatic drainage can lead to the accumulation of inflammatory cells, such as macrophages and neutrophils, within the aortic wall, thereby perpetuating inflammation and tissue destruction^18^. This suggests that enhancing lymphatic function and promoting lymphangiogenesis could be therapeutic strategies to mitigate inflammation and protect the aortic wall from further degradation.

Eosinophils, a type of granulocytic immune cell traditionally associated with allergic responses and parasitic infections, have recently been implicated in cardiovascular diseases, including AAA^12^. Eosinophils are capable of releasing pro-inflammatory cytokines, cytotoxic granules, and reactive oxygen species, which can contribute to tissue damage in the aortic wall^19^. However, eosinophils also play a role in tissue repair and the resolution of inflammation, highlighting their dual nature in disease pathophysiology. Importantly, blood EOS count remains higher in AAA patients than normal controls, and are considered as risk factor of human AAA^11^. Importantly, eosinophils can influence lymphangiogenesis and lymphatic function, thereby modulating the resolution of inflammation^20^. The interaction between eosinophils and the lymphatic system could therefore have significant implications for AAA progression, with the potential to either exacerbate or ameliorate vascular inflammation depending on the balance between their pro-inflammatory and pro-resolving functions^21,22^. The resolution of inflammation is a critical, yet often overlooked, component of immune regulation in AAA. In chronic inflammatory diseases such as AAA, the failure to effectively resolve inflammation can lead to ongoing tissue damage and disease progression^23^. Key to the resolution of inflammation is the clearance of inflammatory cells and the restoration of tissue homeostasis^24^. The lymphatic system plays a pivotal role in this process by facilitating the drainage of immune cells and inflammatory mediators from the aortic wall, thus promoting tissue repair^15^. Enhancing lymphatic function, as well as understanding how eosinophils contribute to both the initiation and resolution of inflammation, could reveal new therapeutic targets aimed at stabilizing the aortic wall and preventing aneurysm rupture.

The current study reports that lymphangiogenesis in AAA control EOS trafficking in aorta wall. This is dependent on specialized pro-resolving mediators (SPM) generated by lymphatic 15LO that regulated the drainage of EOS by the lymphatic system. In summary, the lymphatic system, eosinophils, and the resolution of inflammation are emerging as critical factors in the pathogenesis of AAA. Understanding the interplay between these systems could provide valuable insights into novel therapeutic strategies aimed at modulating immune responses, promoting tissue repair, and ultimately halting the progression of AAA.

## Methods

### Human tissue specimen

From January 2021 to December 2023, 72 patients agreed to be included in the lympheurysm study (Ref CHU: 21163C, Ref INSERM: DIR-20210326016). This inclusion was proposed to patients undergoing open AAA surgery at Toulouse University Hospital (Additional table). The study consisted of taking a sample from the wall of the AAA during the procedure with immediate freezing in liquid nitrogen. The samples included a segment of complete AAA wall, a segment of the so-called inner layer (intima and media) and a segment of the outer layer (adventive). When there was a sufficiently long aneurysmal neck between the renal arteries and the start of the aneurysm (FIG1), a segment of healthy aorta was then taken to serve as a control without modifying the type of intervention or compromising the proximal anastomosis site.

### Lipidomic analysis of human aorta

Lipids corresponding to 10-50 mg of aorta control or intima+media AAA and adventitia were extracted and analysis of bioactive lipids were performed. The extraction protocol and LC-MS/MS analysis were performed by AMBIOTIS SAS (Toulouse, France) using Standard Operating Procedures adapted from Le Faouder *et al.*^30^. Briefly, tissues were dilacerated with scalpel, precisely weighed (about 200 mg) and then crushed with steel beads in methanol (MeOH) using Precellys24 (Bertin Instruments). After protein precipitation and centrifugation, the SPM were extracted from the clear supernatant using HLB Oasis 96 wells extraction plate (Waters). The lipids were finally eluted with methylformate (MeFor) and MeOH. After evaporation of the solvant under N2, the residues were recovered in MeOH/H2O and subjected to LC/MS analysis. Analysis was conducted using a schedule Multiple Reaction Monitoring mode on a 6500+ QTRAP (Sciex) mass spectrometer equipped with an electrospray ionization source in negative mode. The sample was injected bef{Citation}orehand into the Exion LCAD U-HPLC system (Sciex) and eluted on a KINETEX C18 column (2.1*100 mm; 1.7 µm) with a gradient at 0,5 mL/min of buffer A (H2O, 0,1% formic acid (FA)) and buffer B (MeOH, 0,1% FA) placed into a thermostatic oven at 50 °C.

### Quantitative real-time RT-PCR

Total cellular RNA was isolated from human aorta using RNAqueous^®^-Micro Kit (Ambion, USA) according to the manufacturer’s instructions. A total of 1 ug RNA was used to synthesize cDNA using SuperScript® VILO cDNA Synthesis Kit (Ambion, USA). The expression of VEGFR3, Podoplanin, VEGF-C, VEGF-D, IL6, and CCL21 was investigated by SYBR Green real-time reverse transcribed polymerase chain reaction using the ABI StepOne+ Real-time PCR System (Applied Biosystems, Villebon s/ Yvette, France). Each reaction was run with 18S as a reference gene and all data were normalized based on the expression levels of 18S.

hVEGFR-3 F5’- GGA CTC CTG GAC GGC CT; R5’- GGTGTCGATGACGTGTGACT; hPDPN F5’- TGG TTA TGC GAA AAA TGT CGG G; R5’- AGT GTT CCA CGG GTC ATC TT; VEGF-C F5’- AAA GAA GTT CCA CCA CCA AAC; R5’- AGG GAC ACA ACG ACA CAC TTC; hVEGF-D F5’- TGC TGG AAC AGA AGA CCA CTC; R5’-ACA GAC ACA CTC GCA ACG AT; hIL6 F5’- TGC AAT AAC CAC CCC TGA CC; R5’- AGC TGC GCA GAA TGA GAT GA; hCCL21 5’F- CCT TGC CAC ACT CTT TCT CCC; R5’- CAA GGA AGA GGT GGG GTG TA.

#### Mouse model of AAA

All studies received local ethics review board approval and were performed in accordance with the guidelines of the European Convention for the Protection of Vertebrate Animals used for experimentation and according to the INSERM IACUC (France) guidelines for laboratory animal husbandry. All animal experiments were approved by the local branch Inserm Rangueil-Purpan of the Midi-Pyrénées ethics committee, France. Animals from different cages in the same experimental group were selected to assure randomization. Mice were housed in individually ventilated cages in a temperature and light regulated room in a SPF facility and received food and water *ad libitum*. Male C57BL/6J (6 weeks old) were obtained from Envigo, France.

Prox1CreERT2; Alox15^fl/fl^ mice were generated by crossing Prox1CreERT2 mice from Dr Makinen’s laboratory and ALOX15fl/fl obtained from Jackson laboratory (B6.Cg-Alox15^tm1.1Nadl^/J). *12/15-LO^loxP/loxP^* mice possess *loxP* sites flanking exons 2-5 of the arachidonate 15-lipoxygenase (*Alox15*) gene.

#### Mouse model of AAA

10 µl of elastase (Sigma E1250-50MG elastase from porcine pancreas) were applied onto a compress Pangen sponge (Urgo healthcare® laboratories France), which was then placed around the perimeter of the aorta. After 3 weeks, the aorta was harvested and embedded in an OCT block, then frozen. Cryostat sections of 10 µm thickness were prepared for further analysis.

#### Lentiviral injection

The day of the surgery, vehicle or 15-LO expressing lentiviral vector was injected intraperitoneally in a volume of 100μL (10^6^TU/mL).

### Histological analysis of human AAA

AAA was collected at the time of surgery and fixed in 4% formaldehyde in PBS for 1 hour on ice before embedding in paraffin.

Paraffin embedded tissue sections were colored using Masson’s trichrome kit (Biognost), Hematoxylin & Eosin kit (Biognost) and red syrius (Diagnostic Biosystems) following manufacturer’s instructions. Images were acquired on Leica DFC450C camera. Skin thickness was assessed by analysis using Fiji.

### Chemical and reagents

Mouse aorta samples were frozen immediately in O.C.T. Compound (Cellpath) to be stored at −80◦C until the use. Cryosections (5 μm thick) were prepared using CryoSTAP NX50.Frozen tissue section were fixed in ice cold acetone for 2 min then left to dry. They were permeabilized with 0,1% triton, then saturated with 5% BSA in PBS.Samples were incubated with primary antibodies. Rabbit anti-mouse LYVE-1 antibody was from Fitzgerald (Fitzgerald, 70R-LR004 1/200). Anti-mouse eosinophil CD170 (Siglec F) was from Thermo Fisher Scientific (14-1702-82, 1/100).

### In vitro knock down of 15LO

ON-TARGETplus Human ALOX15 siRNA, a guaranteed gene silencing Patented modifications to reduce off-targets, SMARTpool format, and control scramble siRNA were from Dharmacon. Proteins were extracted from HDLEC lysed in RIPA buffer and quantified using the BCA assay. The knockdown of 15-LO was validated by capillary Western blot (JESS) using an anti-human 15-LOX-1 antibody (Abcam, ab119774).

### Transendothelial cell migration assay

Lymphatic transendothelial migration were cultured in Boyden chamber. Eol1 cells (150.000 per well) were incubated on top of the confluent monolayer of HDLEC for 4 hours in the presence of CCL11 (10ng/mL). Filters were harvested and cells that passed through the filter were quantified. Statistical significance was determined using Student’s *t* test.

### Adhesion assay

Eol1 cells were labeled with CellTracker Orange (CMTMR) according to manufacturer’s directions (Invitrogen) and incubated with monolayers of HDLECs in EBM2-culture medium for 20 minutes at 37°C. Cell layers were gently washed with warmed culture medium and fixed in 3.7% 8araformaldehyde prior to enumeration of bound cells per microscopic field at ×200 magnification. Experiments were performed 3 times; results from representative experiments are shown. Statistical significance was determined using Student’s *t* test.

### Statistical analysis

All results presented in this study are representative of at least three independent experiments. Data are shown as the mean ± standard error of the mean (s.e.m.). Statistical significance was determined by two-tailed Student’s *t* test, two-tailed Mann Whitney test, one-way or two-way ANOVA with Tukey post hoc test using Prism ver. 6.0 (GraphPad). Differences were considered statistically significant with a *P* value < 0.05

## Results

### Lymphangiogenesis develops in aorta wall during AAA

To study the lymphangiogenesis development in AAA, histological analysis was performed. Aneurysm aorta was studied in the dilated area and compared to non-dilated area (Fig. 1a). Fibrosis was observed using Sirius red and Masson’s trichrome staining to confirm collagen deposits in aneurysm aorta (Fig. 1b). Then, lymphangiogenesis was evaluated using Lyve-1 immunodetection (Fig. 1c). We observed a significant increase in lymphatic capillary density in the aortic wall compared to control aorta (Fig. 1c and d). Interestingly, lymphatic vessels were located in the adventitia (Fig. 1c). To identify whether the lymphangiogenesis was associated with local gene expression changes, mRNAs were extracted and gene expression analyses were performed by RT-qPCR (Fig. 1e-j). As expected, lymphangiogenesis was associated with an increase in lymphatic endothelial markers VEGFR3 (Fig. 1e) and podoplanin (Fig. 1f). Vascular endothelial lymphatic growth factors C and D (VEGF-C and -D) were slightly increased (Fig. 1g and h). However, we observed a strong induction of pro-inflammatory cytokines such as IL6 (Fig. 1i) and CCL21 (Fig. 1j).

**Figure 1.**
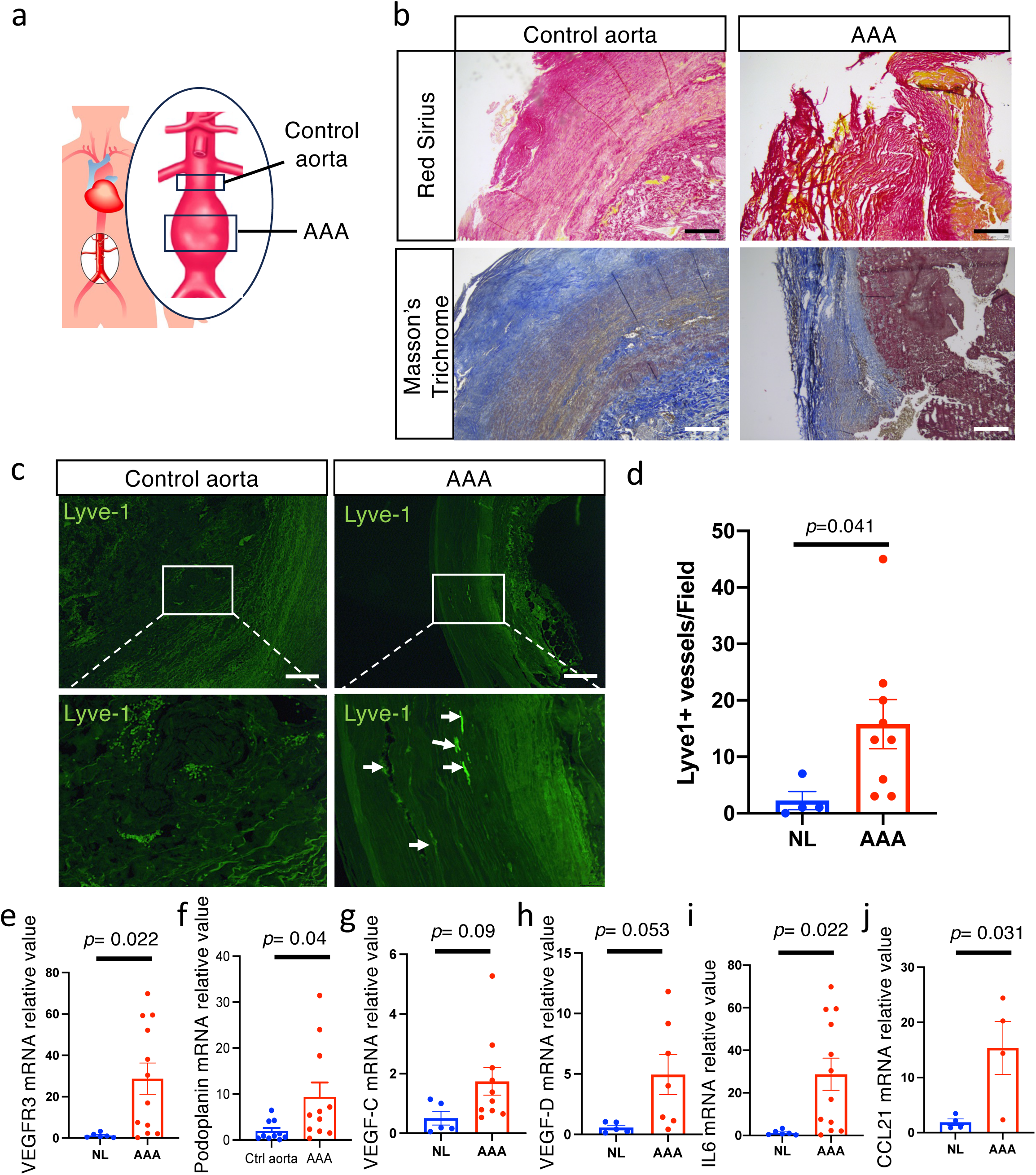
Lymphangiogenesis develops in aorta wall during AAA. **a**, Schematic representation of human aorta showing. **b**, Representative photographs of Hematoxylin-Eosin (H&E, upper panel) and Masson’s trichrome (lower panel) staining. **c**, Aorta wall immunodetection of the lymphatic vessels in AAA. **d**, Quantification of lymphatic vessel density in AAA. **e-j**, VEGFR3 (**e**), Podoplanin (**f**), VEGF-C (**g**), VEGF-D (**h**), IL6 (**i**) and CCL21 (**j**) mRNA relative expression in AAA and control aorta. Scale bars, 100 μm (**a** and **c**).

### Aorta shows decrease in 15LO-derived SPM associated with resolution of inflammation in AAA

Aortic aneurysm occurs when the wall of the aorta weakens and bulges outward due to elastic fibers fragmentation, particularly in the media, which normally provides the aorta with strength and elasticity^25^. This condition is also associated with chronic inflammation within the aortic wall. In that context, several studies have found that patients with AAA exhibit altered levels of specialized proresolving lipid mediators (SPM)^26,14,27,28^.

To understand whether SPM composition could be associated with changes in immune cell recruitment, we performed lipidomic analysis of aneurysmal aorta compared to non-dilated area from the same patient (Fig. 2a, Extended Data Fig. 1). As lymphatic develop predominantly in the adventitia, intima and media were dissected to perform differential analysis (Fig. 2b). We first compared control aorta to AAA. We found an overall downregulation of SPM in AAA without selectivity between Arachidonic Acid (AA), Docosahexaenoic Acid (DHA), and Eicosapentaenoic Acid (EPA) pathways (Extended Data Fig. 1a-f). Among the highly expressed SPM, 15-HETE and 17-HDOHE, two intermediates generated by 15-lipoxygenase (15LO), were significantly downregulated (Fig.2b). We also observed a significant downregulation of 15LO-derived SPM in medium and low-expressed groups including 15HEPE, LxA4, and PD1 (Fig. 2b, Extended Data Fig. 1g). When comparing control aorta to intima/media or to adventitia separately, we found significant decrease in 15HETE in both compartments, whereas 17HDOHE was only significantly decreased in intima-media suggesting that lymphangiogenesis into the adventitia may have an impact on media (Fig. 2c). Similar observations were performed with PDX (Fig. 2d).

**Figure 2.**
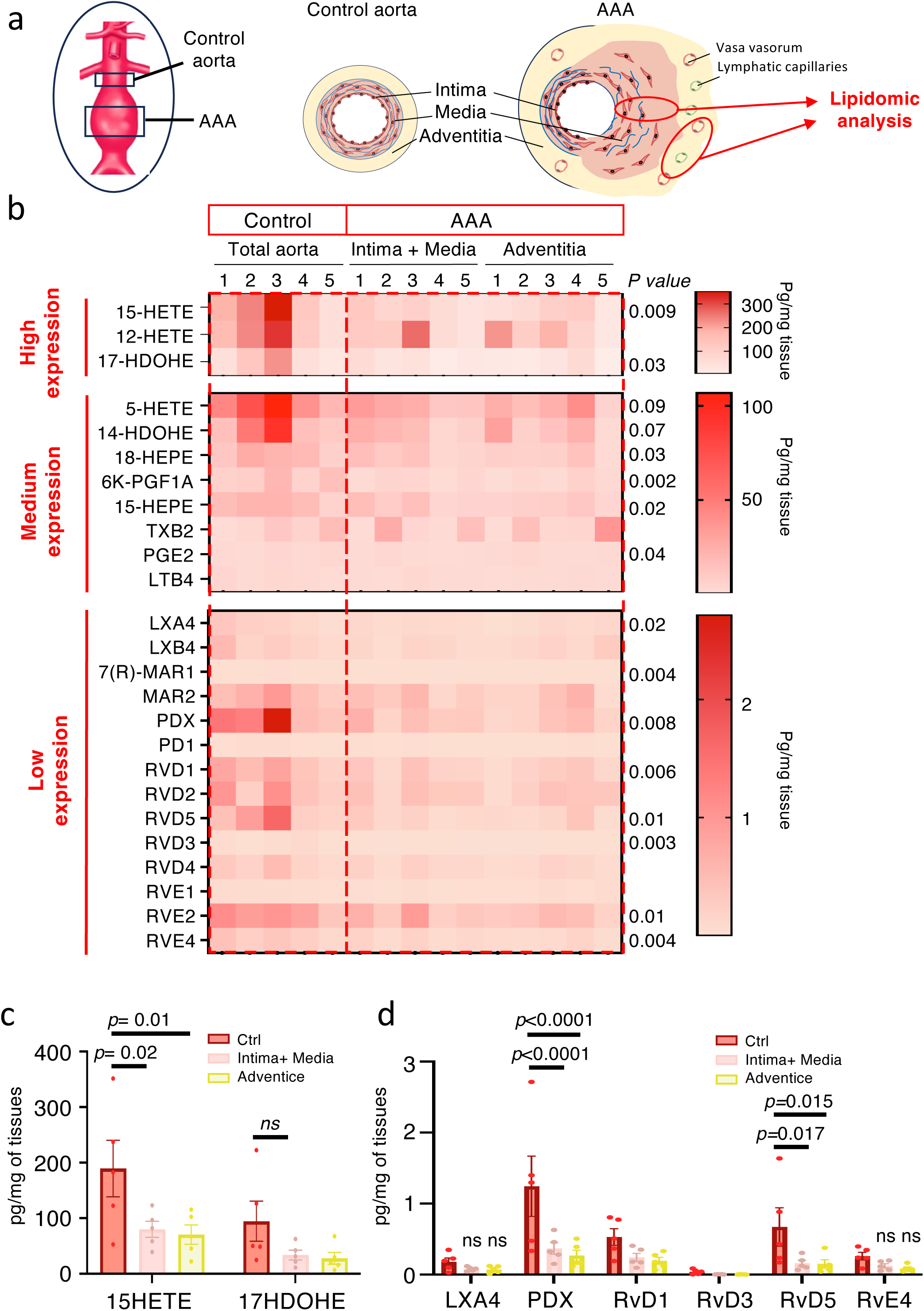
Aorta shows decrease in 15LO-derived SPM associated with resolution of inflammation in AAA. **a**, Schematic representation of the experimental procedure aiming at dissociating intima-media from adventitia before lipidomic chromatography analysis. **b**, Heatmap of lipid mediators showing high, medium and low expression of eicosanoid acids in AAA (pg/mg tissue). **c**, Quantification on intima + media vs adventitia of 15HETE and 17HDOHE, respectively, AA- and DHA-derived lipid mediators generated by 15-LO conversion. **d**, Quantification on intima + media vs adventitia of LXA4, PDX, RvD1, RvD3, RvD5, RvE4 generated by 15-LO conversion.

### Mouse models of AAA show increased lymphangiogenesis in aortic wall

To understand if the lymphatic system had an impact on aortic aneurysm, we studied lymphangiogenesis in two mouse models of AAA (Extended Data Fig. 2 and 3). We first used an established mouse model that promotes AAA three weeks after intraluminal elastase injection into the abdominal aorta (Extended Data Fig. 2a-c). We observed lymphatic vessels development into the adventitia starting 5 days after elastase perfusion (Extended Data Fig. 2d and e). Due to the high mortality associated to this model, we performed another model in which elastase is filed on the external part of the aorta using a loaded sponge (Extended Data Fig. 3a). In this model, AAA develops within 3 weeks (Extended Data Fig. 3a-d). After 21 days, we observed elastic fibers fragmentation associated with aortic wall thickening (Extended Data Fig. 3b), and collagen deposition (Extended Data Fig. 3c and d). As observed in human tissue samples and in the model of elastase perfusion, lymphangiogenesis developed in the adventitia and significantly increased after 14 days (Extended Data Fig. 3e and f).

### Defect in lymphatic endothelial 15LO does not affect aneurysm, by induces increase lymphangiogenesis in aortic wall

Our group recently identified that a defect in lymphatic endothelial 15LO was involved in chronic inflammation associated with a lymphatic disease called lymphedema^15^. To study if lymphatic endothelial 15LO plays a role in the aortic wall inflammation during AAA, we induced AAA in mice selectively depleted for 15LO into the lymphatic endothelium (Alox15^LECKO^ mice)(Fig. 3)^15^. AAA was induced using elastase-loaded sponge assay during 21 days (Fig. 3a). H&E coloration showed elastic fiber fragmentation and aorta wall thickening. However, no difference was found in Alox15^lecKO^ mice compared to control Cre-littermates (Fig. 3b and c). Next, periaortic lymphangiogenesis was assessed using Lyve-1 immunodetection (Fig. 3d). Lymphatic vessel density significantly increased in Alox15^lecKO^ mice (Fig. 3e).

**Figure 3.**
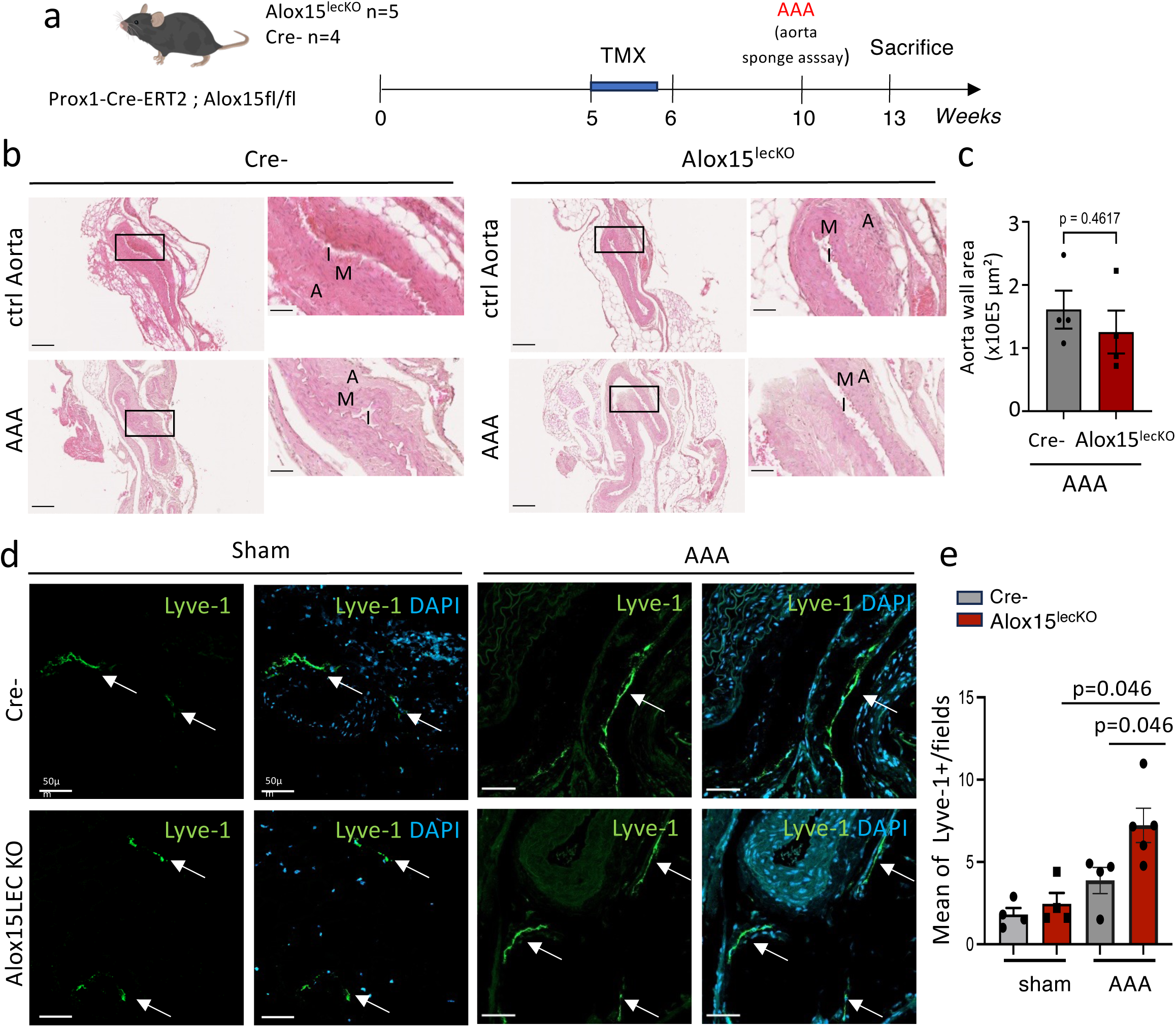
Defect in lymphatic endothelial 15LO does not affect aneurysm, by induces increase in aortic wall lymphangiogenesis. **a**, Schematic representation of the experimental procedure: 15LO lymphatic depletion is induced by tamoxifen (TMX) gavage during 5 days. 5 weeks of TMX wash out are performed before proceeding to AAA induction. **b**, Representative images of Hematoxylin-Eosin (H&E) aorta staining in Alox15^lecKO^ mice and control littermates. **c**, Quantification of aorta wall area. **d**, Aorta wall immunodetection of the lymphatic vessels in AAA from Alox15^lecKO^ mice and control littermates. **e**, Quantification of lymphatic vessel density in AAA. Scale bars, 200 μm (**b-c** left panel), 50 μm (**b-c** right panel and **d**).

### Defect in lymphatic endothelial 15LO induces increase in eosinophil cell population

To evaluate whether a difference in periaortic lymphatic vessels density could affect immune cells trafficking, we performed systemic blood count and tissue immunodetection (Fig. 4). Increase in adventitia lymphangiogenesis was not associated with significant changes in circulating immune cell populations including neutrophils (Fig. 4a), lymphocytes (Fig. 4b), and monocytes (Fig. 4c). Surprisingly, systemic blood eosinophil count was significantly upregulated in Alox15^lecKO^ mice compared to control Cre-littermates (Fig. 4d).

**Figure 4.**
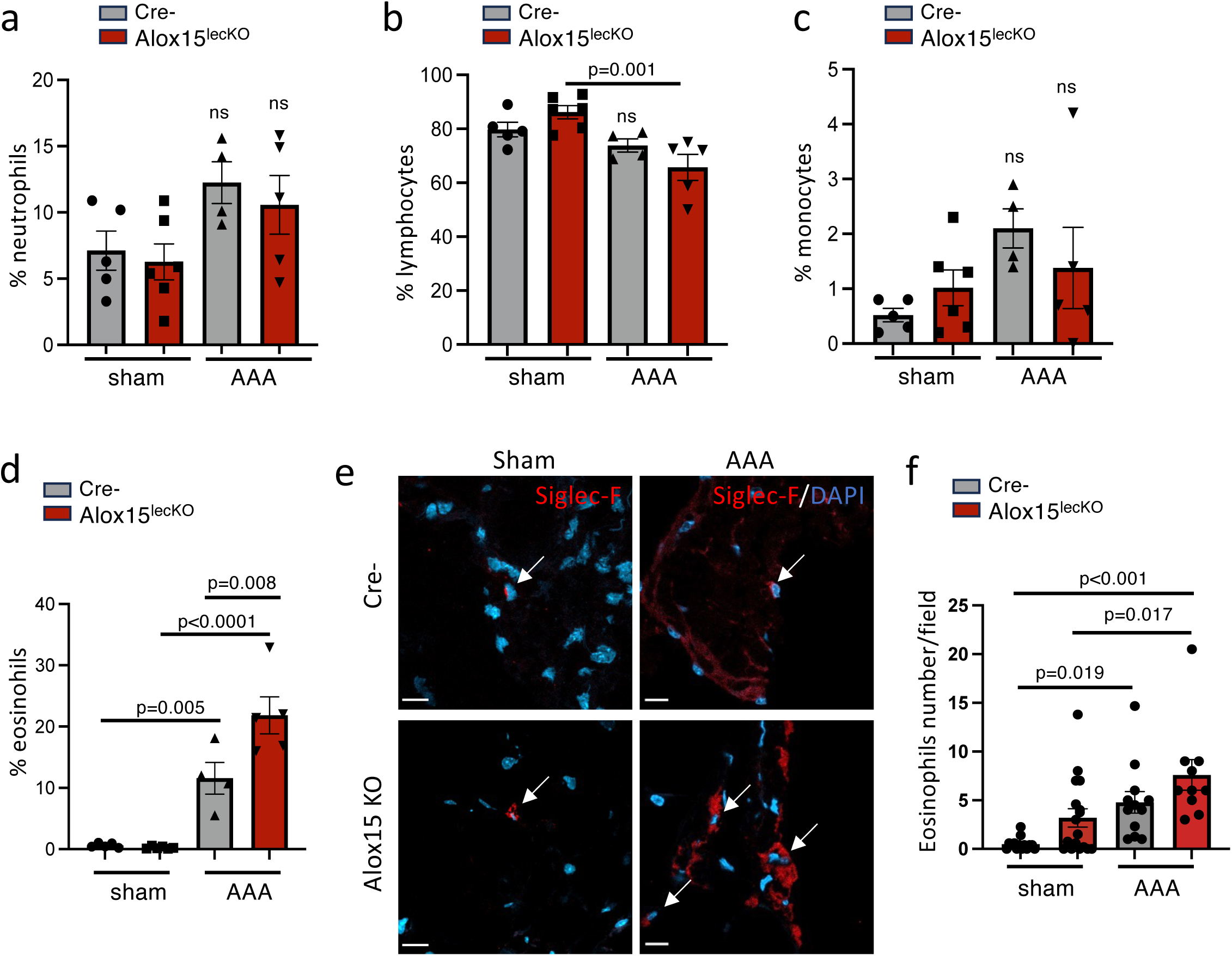
Defect in lymphatic endothelial 15LO induces increase in eosinophil cell population. **a-d,** AAA-related alterations in white blood cell differential count: neutrophil (a), lymphocyte (b), monocytes (c), and eosinophils (d). **e**, Aorta wall immunodetection of eosinophils in AAA from Alox15^lecKO^ mice and control littermates (Cre-). **f**, Quantification of eosinophil cells density in AAA. Scale bars, 10 μm (**e**).

We next studied eosinophils infiltration into the aorta wall (Fig. 4e and f). An immunodetection showed a significant increase in eosinophil cell count in the aorta wall from AAA mice compared to control sham (Fig. 4e and f).

### Loss of lymphatic endothelial 15LO *in vitro* increases transendothelial migration

We next investigated *in vitro* whether lymphatic endothelial 15LO could modify eosinophil numbers in aorta wall (Fig. 5). Knock-down of 15LO was performed in human dermal lymphatic endothelial cells (HDLEC) using smart pool siRNA (Fig. 5a and b). Knock-down was confirmed by western blot (Fig. 5a and b). We first studied the adhesion of eosinophils on HDLEC (Fig. 5c and d). The knockdown of 15LO in HDLEC significantly reduced the eosinophil adhesion to the lymphatic endothelial monolayer (Fig. 5d). Next, we evaluated the lymphatic transendothelial migration of eosinophils in boyden chambers (Fig. 5e and f). The eosinophil migration was stimulated by CCL11, the main chemotactic mediator responsible for the migration of eosinophils. We observed a significant induction of lymphatic transendothelial migration of eosinophil in 15LO knockdown HDLEC (Fig. 5e and f). These results suggest that lymphatic 15LO regulates the drainage of tissular eosinophils across the lymphatic system to make them join the bloodstream.

**Figure 5.**
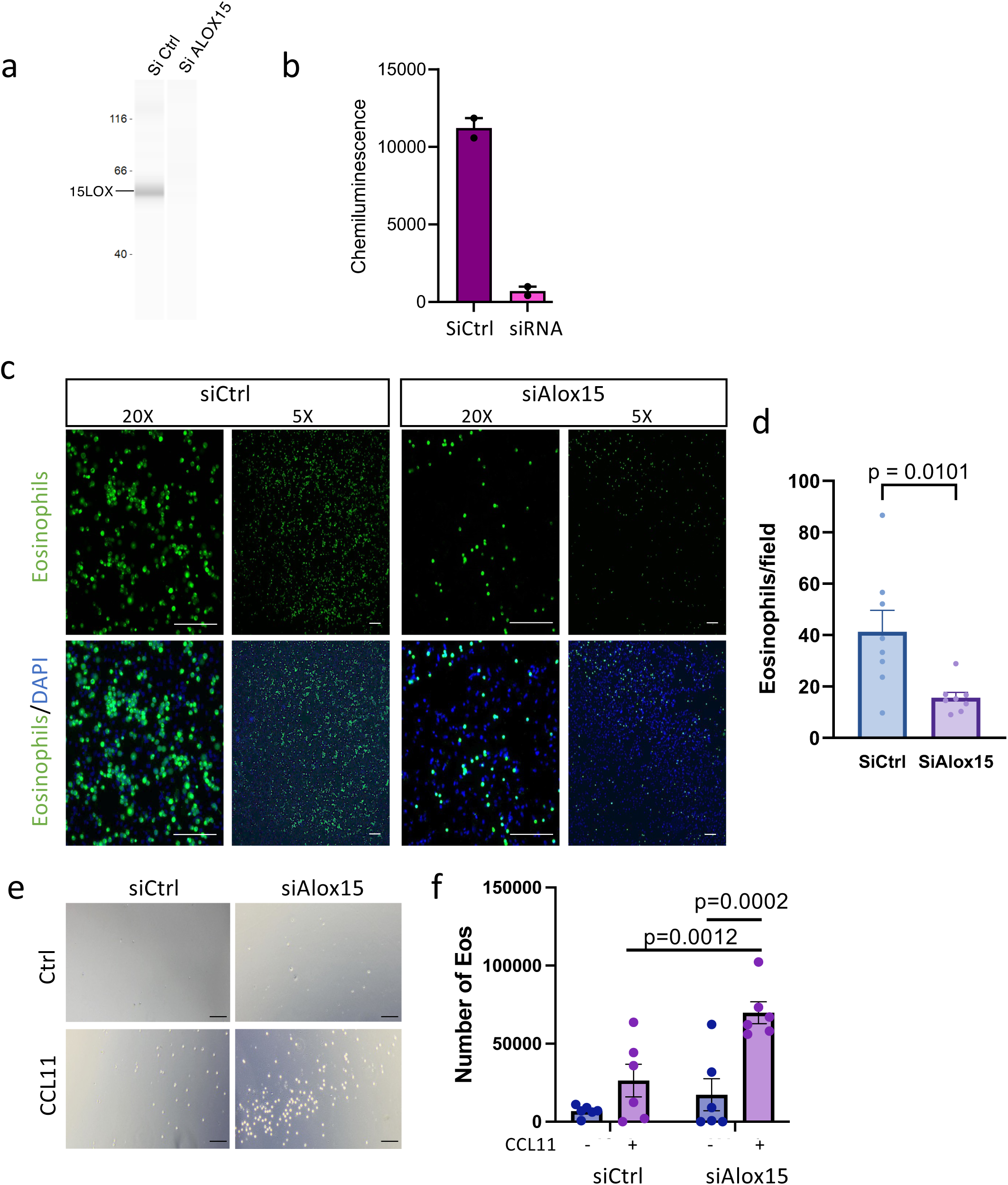
Loss of lymphatic endothelial 15LO *in vitro* increases transendothelial migration. **a**, Representative western blot of the siRNA transfection experiment in HDLEC. **b**, Graph showing a significant reduction of the 15LO protein level in the silenced cells, as compared to the negative Si scramble (SiCtrl) control. **c**, Fluorescent micrographs of eosinophil adhesion to lymphatic endothelial cells depleted for ALOX15. **d**, Quantification of the number of adherent eosinophils per field. **e**, Micrographs of eosinophil after lymphatic endothelial transmigration. **f**, Quantification of the number of migrating eosinophils per field. Scale bars 50 μm (**c**), 100 μm (**e**).

### 15LO rescue increase the resident eosinophils in AAA to reduce the lesion

To confirm the impact of the loss of 15LO-derived SPMs in AAA, we generated a lentivector (LV) that allows stable overexpression of 15LO (LV15LO) after transduction (Fig. 6a). LV15LO was injected intraperitoneally at the time of surgery. A control lentivector (LV Ctrl) was also used as a control. As shown in Fig.4, number of circulating eosinophils was increased in AAA mice (Fig. 6b). We observe a slight reduction of eosinophils blood count in Cre-mice during AAA. Importantly, we found a significant reduction of blood circulating eosinophils in Alox15^lecKO^ mice (Fig. 6b). The lymphatic endothelial knockdown of 15LO had no effect on neutrophil and lymphocyte blood counts (Extended Data Fig. 4a-b).

**Figure 6.**
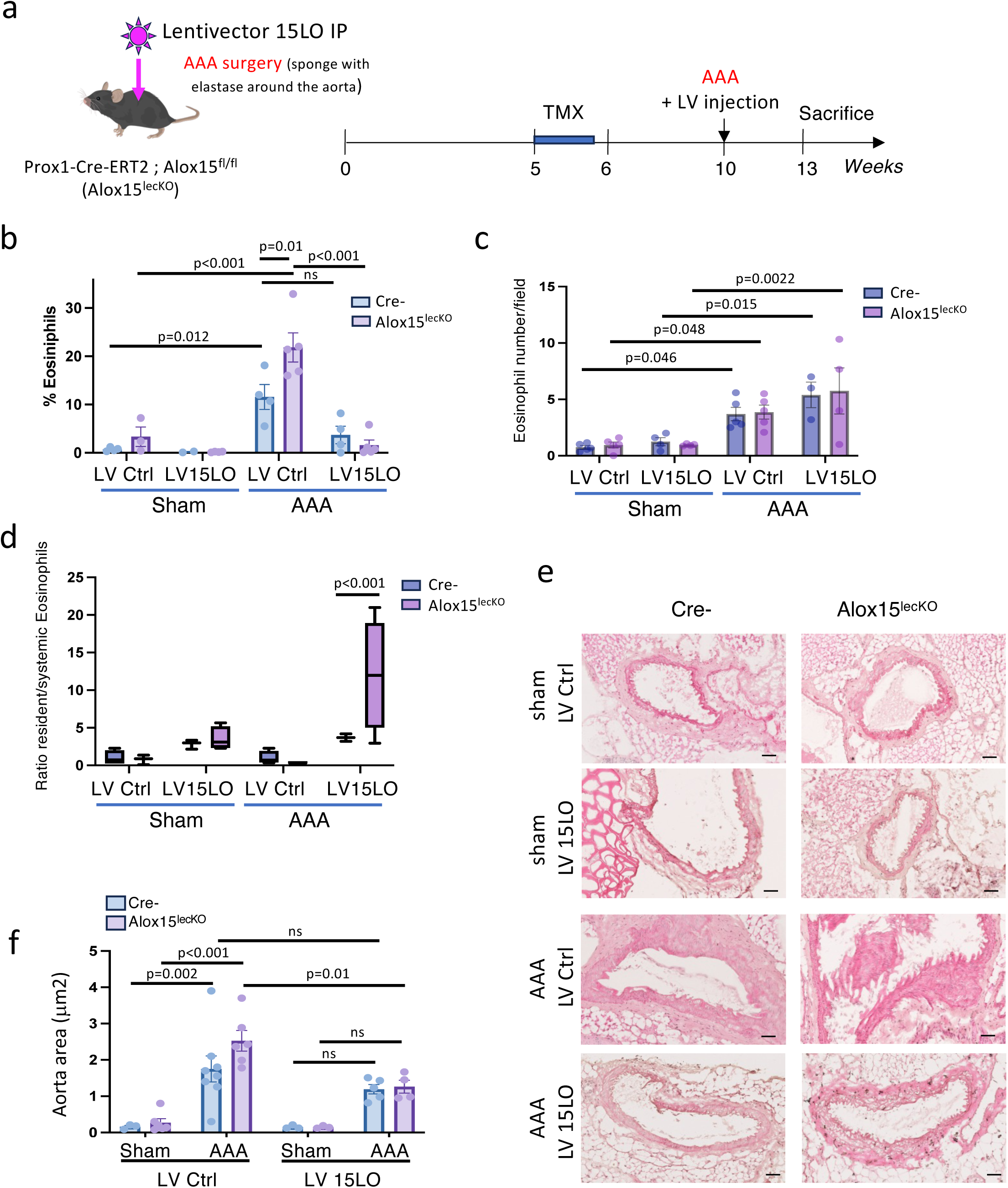
15LO rescue improves the number of resident eosinophils and reduced AAA. **a**, Schematic representation of the experimental procedure: 15LO lymphatic depletion is induced by tamoxifen (TMX) gavage during 5 days. 5 weeks of TMX wash out are performed before proceeding to AAA induction and IP lentiviral injection. **b**,Percentage of blood circulating eosinophils in Alox15^lecKO^ mice injected with 15LO lentivector. **c**, Quantification of eosinophil cells density in AAA from Alox15^lecKO^ mice injected with 15LO lentivector. **d**, Ratio of resident/systemic eosinophils. **e**, Representative images of Hematoxylin-Eosin (H&E) aorta staining from Alox15^lecKO^ mice and control littermates injected with 15LO lentivector. **f**, Quantification of aorta wall area from Alox15^lecKO^ mice and control littermates injected with 15LO lentivector. Scale bars, 50 μm (**e**).

Interestingly LV15LO was also associated with an increase in resident eosinophils in the aneurysmal aorta wall compared to sham (Fig. 6c). However, when comparing the ratio between circulating and resident eosinophils, we found that 15LO rescue significantly increased the eosinophil number in the AAA lesion compared to circulating ones (Fig. 6d). To evaluate whether the decrease in aorta eosinophils was associated with a lack of cell proliferation, KI67 immunodetection was performed (Extended Data Fig. 5a and b). In both Alox15^lecKO^ mice and in control Cre-littermates, no difference in eosinophils proliferation was observed in aneurysmal aorta compared to control aorta suggesting that changes in aorta wall lymphangiogenesis controls eosinophils trafficking rather than proliferation. We next evaluated the effect of 15LO LV on AAA development (Fig. 6e and f). In accordance with the literature showing that aortic eosinophils have a protective effect on AAA, we observed a reduction in the lesion size after LV15LO injection (Fig. 6f).

## Discussion

Our findings underscore the pivotal role of lymphangiogenesis and its regulation in immune cell trafficking, particularly in the context of chronic inflammatory diseases such as abdominal aortic aneurysm (AAA). We observed that the physiologic functions of lymphatic vessels in immune surveillance and immune cell trafficking, which are well-documented in conditions like graft rejection and Crohn’s disease^29^, may also extend to AAA. The disruption of this balance between resident and circulating immune cells in inflammatory diseases highlights a potential mechanism contributing to the pathogenesis of AAA.

In our study, we investigated the role of 15-lipoxygenase (15LO) in the lymphatic system and its impact on immune regulation in AAA. Our previous work in lymphedema demonstrated that a lack of 15LO impairs the resolution of chronic inflammation, suggesting a broader relevance of this enzyme in lymphatic function^15^. Here, we extend these findings to AAA, where a loss of specialized pro-resolving mediators (SPMs) generated by 15LO correlates with changes in lymphatic density in the aortic wall. This reduction in lymphatic density may compromise the immune-modulatory functions of the lymphatic system, exacerbating inflammatory processes in AAA.

To establish causality, we utilized transgenic mice with lymphatic-specific depletion of 15LO. Consistent with previous reports, we observed elevated eosinophil blood counts in AAA^11^. However, our data revealed that lymphatic-specific 15LO depletion exacerbates this increase, indicating an important role of the lymphatic system in regulating eosinophil trafficking. Mechanistically, in vitro assays demonstrated that 15LO depletion in lymphatic endothelial cells reduces eosinophil adhesion to the lymphatic endothelium while increasing their transendothelial migration. These findings suggest that 15LO deficiency promotes the mobilization of eosinophils from tissue to the lymphatic system and subsequently to the bloodstream, thereby reducing the pool of protective eosinophils within the aortic lesion. This depletion of resident eosinophils likely contributes to the exacerbation of inflammation and AAA progression.

Importantly, we demonstrated that rescuing 15LO expression using lentiviral vectors ameliorates these effects. Restoration of 15LO expression reduced both circulating and resident eosinophil counts, with a more pronounced effect on circulating eosinophils, thereby increasing the ratio of resident to circulating eosinophils. This shift in the resident-to-circulating eosinophil ratio was associated with a significant reduction in AAA lesion size. Notably, this effect was independent of cell proliferation, highlighting the central role of eosinophil trafficking and localization rather than their absolute numbers in modulating disease outcomes. Also, the effect observed in Cre-control mice corresponds to the entire set of cells transduced by the lentivector, while the difference observed between Cre- and Alox15^lecKO^ mice can be attributed to lymphatic endothelial 15LO.

These findings suggest a novel paradigm wherein the lymphatic system, via 15LO-mediated regulation, governs the balance between resident and circulating immune cells. The observed impact on eosinophil trafficking and AAA progression emphasizes the importance of maintaining lymphatic homeostasis in inflammatory diseases. Moreover, our data suggest that therapeutic strategies aimed at enhancing lymphatic function or restoring 15LO activity may represent promising avenues for the treatment of AAA and potentially other chronic inflammatory diseases^11^.

Further investigations are warranted to elucidate the broader implications of these findings, including the identification of additional immune cell subsets regulated by the lymphatic system and the exploration of potential cross-talk between lymphatic endothelial cells and other components of the immune system. Understanding the precise molecular mechanisms underlying these interactions may reveal new therapeutic targets to modulate immune responses and improve outcomes in inflammatory diseases.

## Acknowledgements

We thank E. Lhullier, F. Martins (GenoToul platform), Zakarof A., and Riant E. (TRI platform) for their technical support as well as M. Rousseau from the platform Anexplo Genotoul (Inserm US006, Toulouse, France) for their outstanding technical assistance.We thank the imaging Phi platform of I2MC Institute (R. Flores, C. Segura). We thank Taija Makinen for providing Prox1-CreERT2 mice.

## Author contributions

B.G.S. conceived the study. A.H. performed human aorta dissection, conceived the tissue collection, performed the mousee AAA endovascular model and analyzed the data. L. V., E. B., A. B. performed mice experiments and histological analyses. M.D. participated in the design of lipidomic experiments. J. M. D., F. M, E. L., contributed to the experimental design. A. L. and X. C. contributed to the human tissue collection. A.C.P. designed the lentivector expreiment. B.G.S. designed the study, performed data analysis, and wrote the manuscript.

## Supplemental information

### Extended Data Figure legend

**Extended Data Figure 1.**
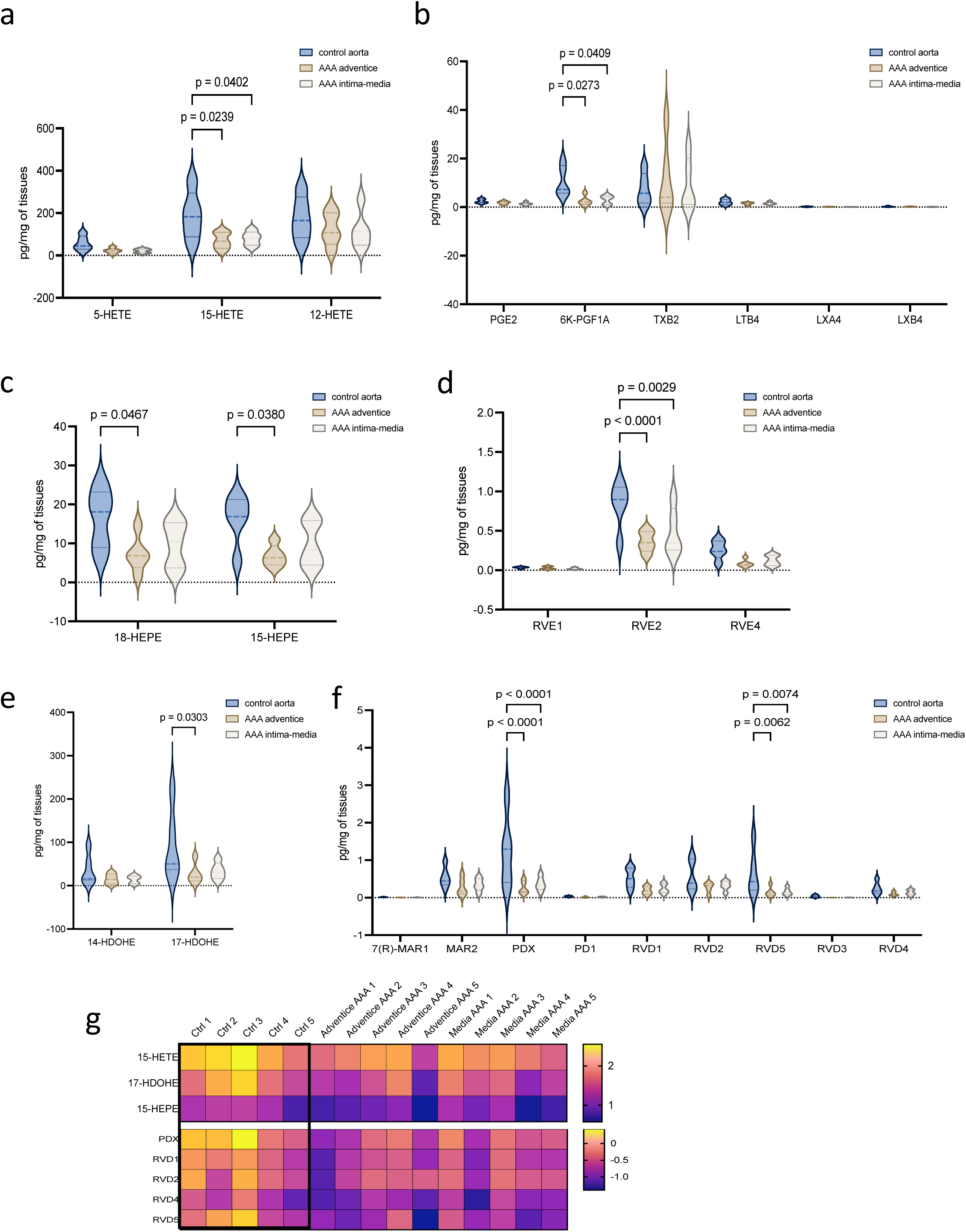
**a**, Quantification of control aorta, AAA intima + media, and AAA adventitia of Arachidonic Acid-derived lipid mediators 5HETE, 15HETE, and 12HETE. **b**, Quantification of control aorta, AAA intima + media, and AAA adventitia of Arachidonic Acid-derived lipid mediators PGE2, 6K-PGF1A, TXB2, LTB4, LXA4, and LXB4. **c**, Quantification of control aorta, AAA intima + media, and AAA adventitia of Eicosapentaenoic Acid-derived lipid mediators 18-HEPE and 15-HEPE. **d**, Quantification of control aorta, AAA intima + media, and AAA adventitia of Eicosapentaenoic Acid-derived lipid mediators RVE1, RVE2, and RVE4. **e**, Quantification of control aorta, AAA intima + media, and AAA adventitia of Docosahexaenoic Acid-derived lipid mediators 14-HDOHE and 17-HDOHE. **f**, Quantification of control aorta, AAA intima + media, and AAA adventitia of Docosahexaenoic Acid-derived lipid mediators 7(R)-MAR1, MAR2, PDX, PD1, RVD1, RVD2, RVD5, RVD4, and RVD4. **g**, Heatmap of lipid mediators generated by 15-LO conversion.

**Extended Data Figure 2.**
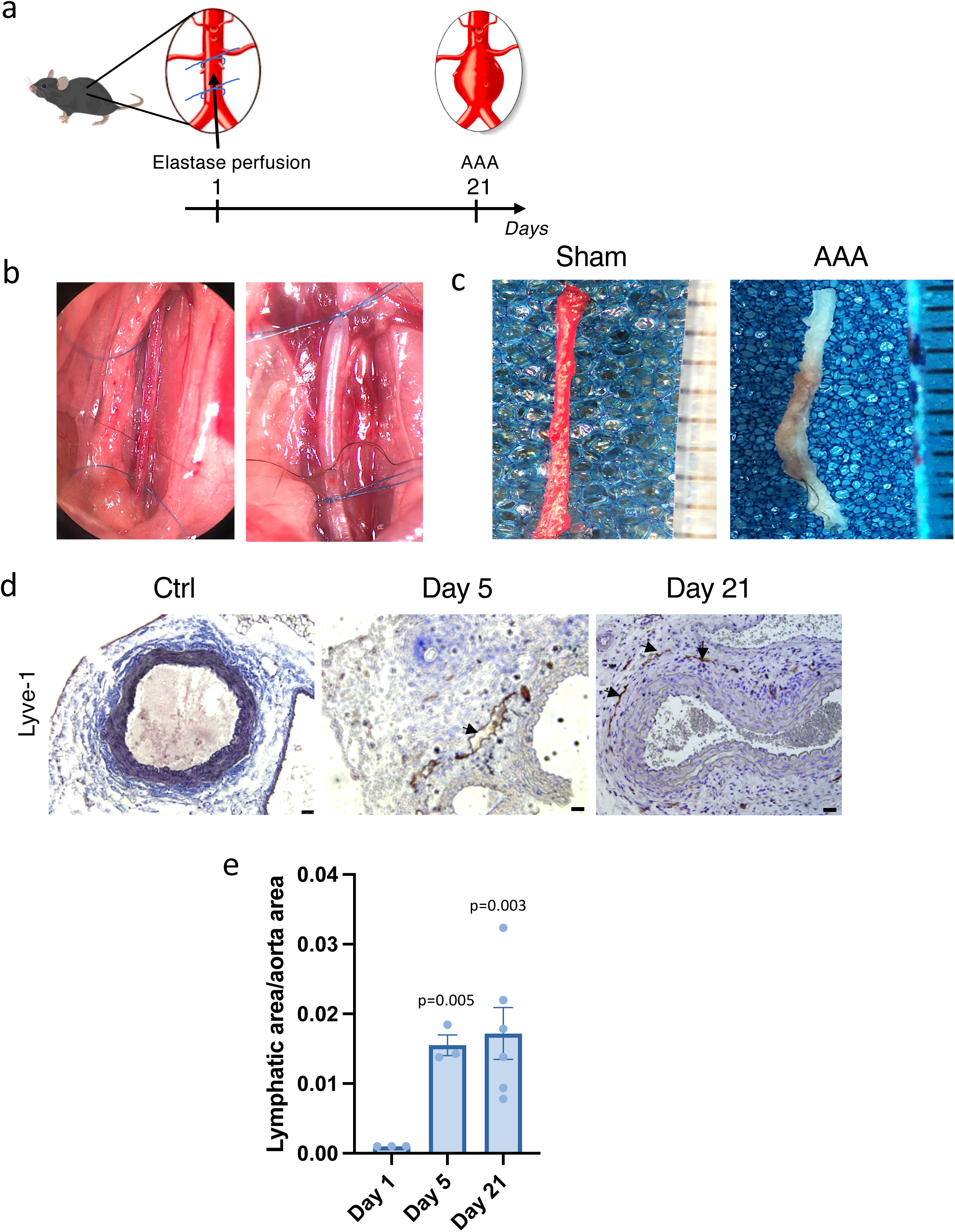
**a**, Schematic representation of the experimental procedure that induces mouse AAA using intraluminal elastase injection. **b**, Representative images of mouse aorta exposed by midline laparotomy. 8.0 polypropylene wires were tied around the aorta to perform proximal and distal clamping. **c**, Representative images of mouse abdominal aorta perfused with saline solution (sham) or elastase. **d**, Aorta wall immunodetection of the lymphatic vessels in AAA. **e**, Quantification of lymphatic vessel area in AAA. Scale bars, 50 μm (**d**).

**Extended Data Figure 3.**
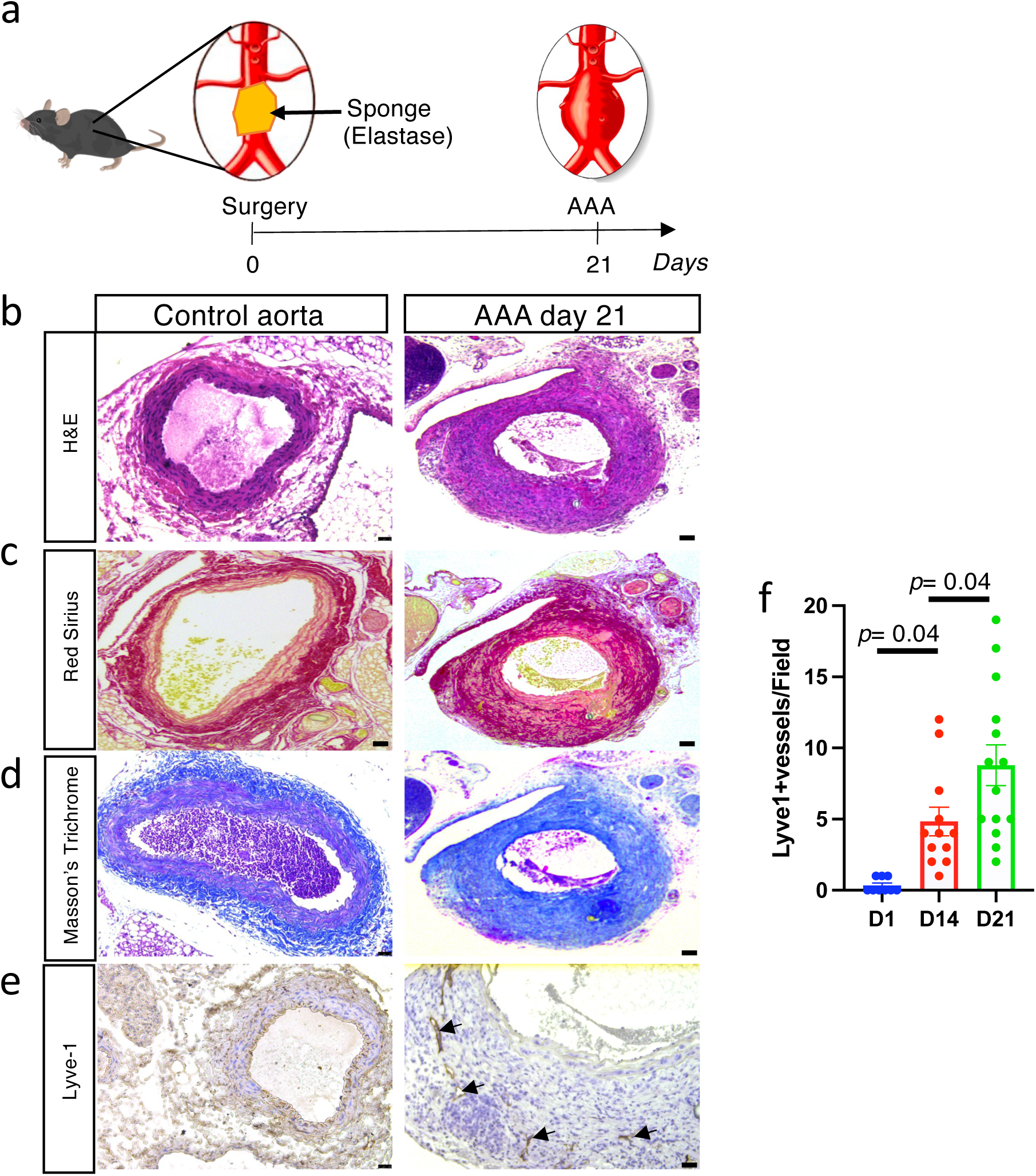
**a**, Schematic representation of the experimental procedure that induces mouse AAA using elastase-containing sponge. **b**, Representative images of Hematoxylin-Eosin (H&E) staining. **c**, Representative images of Red Syrius staining. **d**, Representative images of Masson’s trichrome staining. **e**, Aorta wall immunodetection of the lymphatic vessels in AAA. **f**, Quantification of lymphatic vessel density in AAA. Scale bars, 50 μm (**b-e**).

**Extended Data Figure 4.**
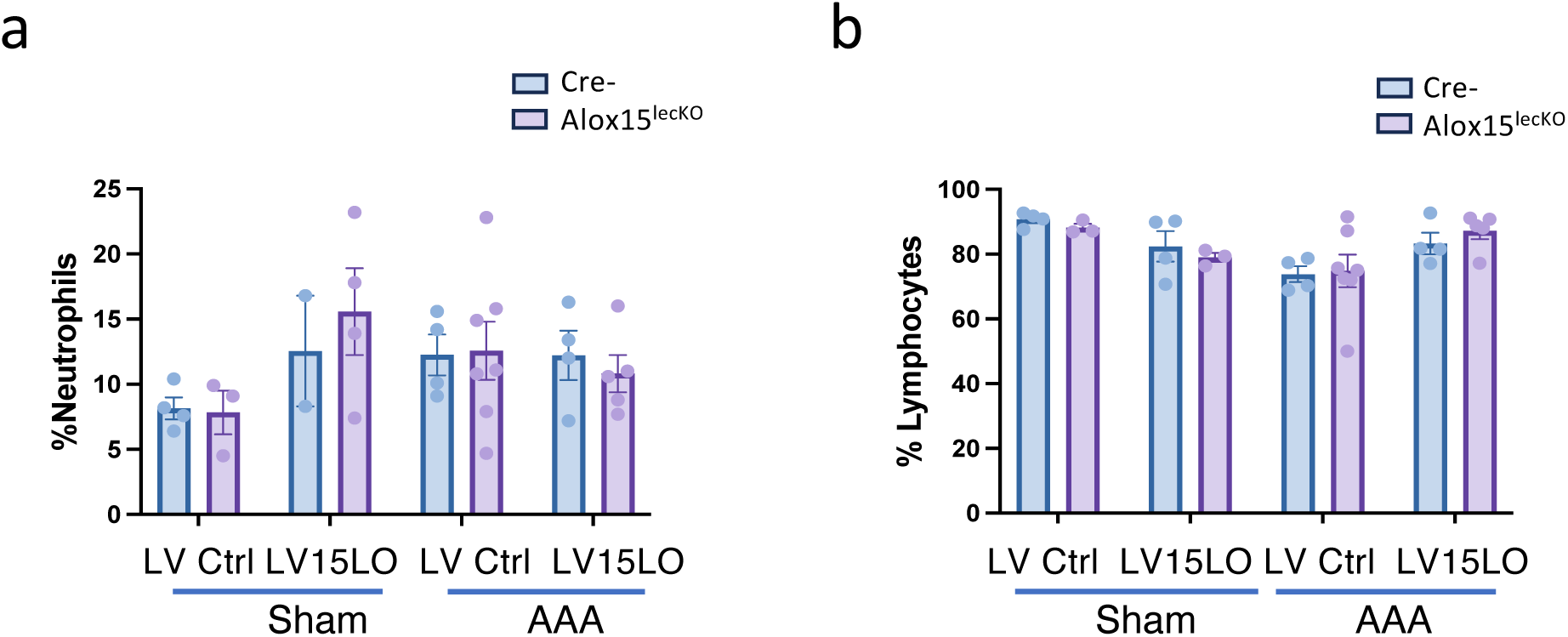
Percentage of blood circulating neutrophil (**a**) and lymphocyte (**b**) in control (sham) and AAA mice injected with 15LO lentivector.

**Extended Data Figure 5.**
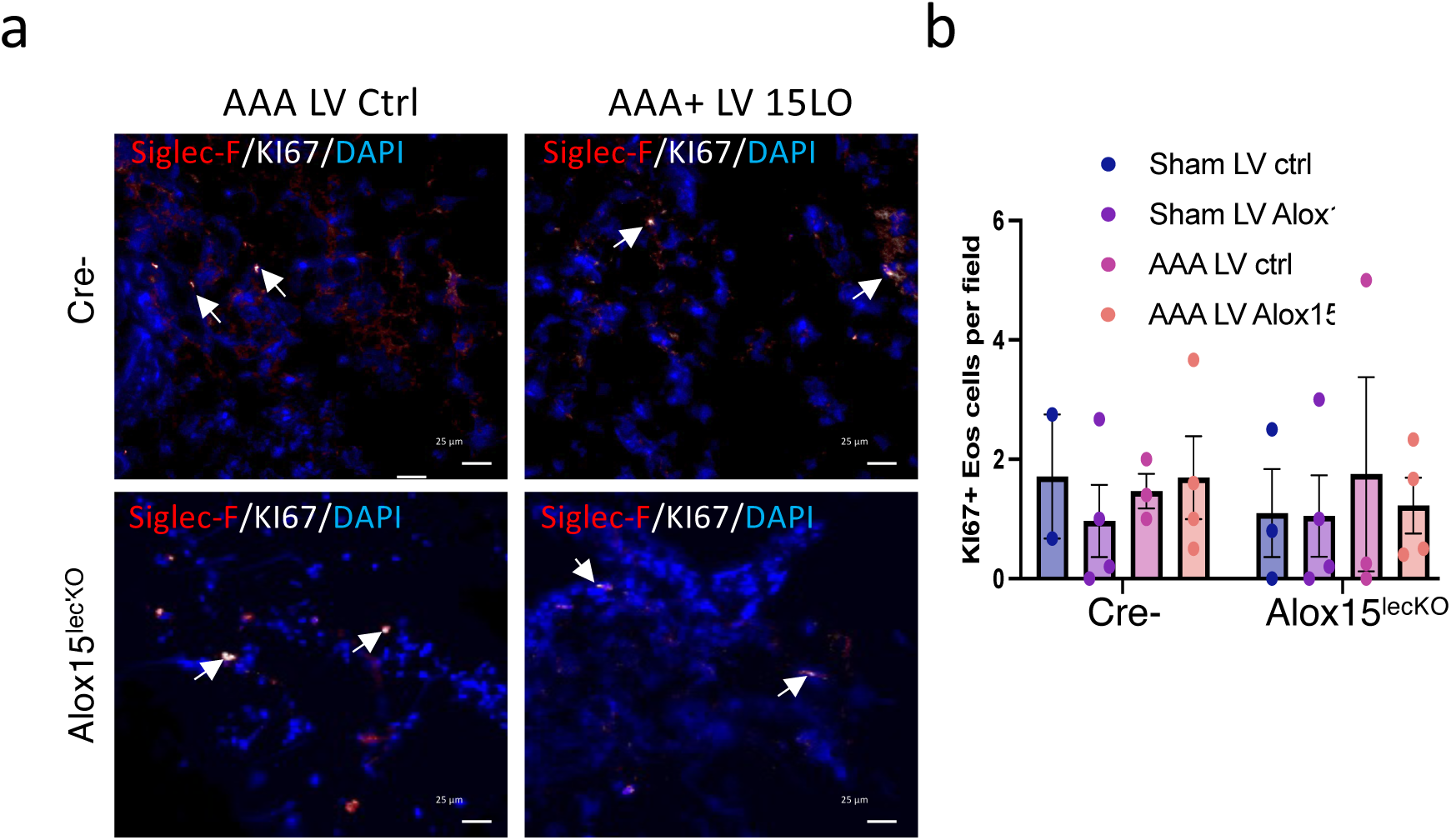
**a**, Aorta wall immunodetection of KI67-positive eosinophils in AAA from Alox15^lecKO^ mice and control littermates (Cre-) injected with 15LO lentivector. **b**, Quantification of KI67+ eosinophil cells density. Scale bars, 10 μm (**d**).

